# Adaptive mutations in RNA polymerase and the transcriptional terminator Rho have similar effects on *Escherichia coli* gene expression

**DOI:** 10.1101/089268

**Authors:** Andrea González-González, Shaun M. Hug, Alejandra Rodríguez-Verdugo, Jagdish Suresh Patel, Brandon S. Gaut

**Affiliations:** Department of Ecology and Evolutionary Biology, University of California, Irvine, CA, USA; Department of Biological Sciences, University of Idaho, Moscow, ID, USA; Department of Environmental Systems Sciences, ETH Zürich, Zürich, Switzerland; Department of Environmental Microbiology, Eawag, Dübendorf, Switzerland; Center for Modeling Complex Interactions, University of Idaho, Moscow, ID, USA

**Keywords:** transcriptional termination, protein structure, relative fitness, experimental evolution, adaptive pathways, phenotypic convergence

## Abstract

Modifications to transcriptional regulators play a major role in adaptation. Here we compared the effects of multiple beneficial mutations within and between *Escherichia coli rpoB*, the gene encoding the RNA polymerase β subunit, and *rho*, which encodes a transcriptional terminator. These two genes have harbored adaptive mutations in numerous *E. coli* evolution experiments but particularly in our previous large-scale thermal stress experiment, where the two genes characterized two alternative adaptive pathways. To compare the effects of beneficial mutations, we engineered four advantageous mutations into each of the two genes and measured their effects on fitness, growth, gene expression and transcriptional termination at 42.2°C. Among the eight mutations, two *rho* mutations had no detectable effect on relative fitness, suggesting they were beneficial only in the context of epistatic interactions. The remaining six mutations had an average relative fitness benefit of ∼20%. The *rpoB* mutations altered the expression of ∼1700 genes; *rho* mutations altered the expression of fewer genes, most of which were a subset of the genes altered by *rpoB*. Across our eight mutants, relative fitness correlated with the degree to which a mutation restored gene expression back to the unstressed, 37.0°C state. The *rho* mutations do not enhance transcriptional termination in known *rho*-terminated regions, but the genome-wide effects of mutations in both genes was to enhance termination. Although beneficial mutations in the two genes did not have identical effects on fitness, growth or gene expression, they acted predominantly through parallel phenotypic effects on gene expression and transcriptional termination.

## INTRODUCTION

In 1975, King and Wilson famously speculated that evolutionary modifications in phenotype “… are more often based on changes in the mechanisms controlling the expression of genes than on sequence changes in proteins” (King and Wilson, 1975). The relative frequency of changes in regulatory vs. structural genes is still debated (Hoekstra and Coyne, 2007, Carroll, 2008, Wittkopp and Kalay, 2011), but it has nonetheless become clear that regulatory genes do often play a crucial role in adaptation. For example, evolutionary modifications of transcriptional regulators have modified plant phenotypes during domestication (Doebley et al., 2006, Olsen and Wendel, 2013), contributed to the shift from marine to freshwater environments for sticklebacks (Peichel and Marques, 2017), and facilitated the adaptation of microbes during laboratory evolution experiments (Conrad et al., 2011, Long et al., 2015). Microbial evolution experiments have further shown that mutations in global transcriptional regulators are often highly beneficial, occur early in the course of adaptation, and appear under different selective regimes (Long et al., 2015). The last observation is especially true in *E. coli* evolution, because experiments over a wide range of strains and growth conditions have consistently identified adaptive genetic changes within the genes that encode the RNA polymerase (RNAP) complex [e.g., (Herring et al., 2006, Conrad et al., 2009, Charusanti et al., 2010, Conrad et al., 2010, Tenaillon et al., 2012, Sandberg et al., 2014, Deatherage et al., 2017)] and the Rho transcriptional terminator [e.g., (Freddolino et al., 2012, Tenaillon et al., 2012, Herron and Doebeli, 2013, Le Gac et al., 2013, Haft et al., 2014, Lee and Helmann, 2014, Dillon et al., 2016, Deatherage et al., 2017)].

These observations raise at least three interesting topics about adaptive changes within *E. coli* transcriptional regulators. The first, which we do not address here, is why modifications of global regulators like RNAP and Rho are favored in evolution experiments, despite the fact that RNAP function and sequence is often highly conserved in nature (Long et al., 2015). The second concerns potential pleiotropic effects, because some transcriptional regulators can, in theory, affect the expression of every gene in the genome. Indeed, single adaptive mutations in the *rpoB* gene, which encodes the RNA Polymerase *β* subunit, alter the expression of hundreds to thousands of downstream genes (Sandberg et al., 2014, Rodríguez-Verdugo et al., 2016, Utrilla et al., 2016). These findings raise the possibility that some of the shifts in gene expression are maladaptive. Consistent with this view, at least two studies have shown that fitness gains after the fixation of a beneficial *rpoB* mutation act to ameliorate negative pleiotropic effects (Sandberg et al., 2014, Rodríguez-Verdugo et al., 2016).

A third topic is about the relative effects of different adaptive mutations, both within the same transcriptional regulator (e.g., different mutations within *rpoB*) and between transcriptional regulators (e.g., *rpoB* vs. *rho*). These are potentially interesting contrasts because it has been shown that different mutations can have different effects on fitness, pleiotropy and evolutionary contingency (Rosenblum et al., 2014), even for different mutations within the same gene (Linnen et al., 2009). It seems likely that there are large-scale differences in the effects of mutations to RNAP and Rho. A modification of RNAP has the potential to affect the expression of every gene, but Rho terminates transcription for a subset of ∼20% to 50% of *E. coli* genes (Cardinale et al., 2008, Peters et al., 2009). This begs the questions: Do adaptive mutations in *rho* affect the expression of fewer genes? If so, are mutations in *rho* generally more beneficial, because they cause fewer negatively pleiotropic effects? And, to what extent are these effects consistent across different beneficial mutations within the same transcriptional regulator?

To our knowledge, the effects of adaptive mutations have not yet been compared explicitly between different transcriptional regulators. This may be due, in part, to the fact that a valid comparison requires that mutations in the two regulatory genes are adaptive in the same genetic background under identical culture conditions. Fortunately, our previous thermal adaptation experiment has yielded appropriate mutations. Our experiment evolved 115 populations of *E. coli* independently from the same ancestral clone (REL1206) for 2000 generations at 42.2°C (Tenaillon et al., 2012). At the end of the experiment, we isolated one clone from each population and found that fitness had improved ∼40%, on average, relative to REL1206. We also sequenced the evolved clones and identified over 1300 mutations. Using parallelism as a *prima facie* argument for adaptation (Wichman et al., 1999, Fong et al., 2005, Stern, 2013), about half of the observed 1300 mutations were adaptive, either because the same mutation occurred in more than one clone or because mutations occurred in the same gene in different clones.

Two of the most mutated genes in our thermal stress experiment were *rpoB*, which accumulated a total of 87 mutations in 76 of the 115 clones, and *rho,* which accumulated 24 different nonsynonymous mutations across 43 of 115 clones. Mutations in the two genes were found together less often than expected at random, suggesting negative epistatic interactions (Tenaillon et al., 2012). More generally, the two genes define alternative pathways to thermal stress adaptation. The first includes mutations in *rpoB*, as well as mutations in the other RNA polymerase subunits (*rpoA*, *rpoC* and *rpoD*) and six *rod* genes that affect cell shape. In contrast, the *rho* pathway includes mutations of the cardiolipin synthase (*cls*) gene and the transcription factor gene *iclR.* We have shown that these two adaptive pathways differ in some aspects of their phenotypes, such as chemical sensitivities (Hug and Gaut, 2015) and trade-offs for growth at low temperature (Rodriguez-Verdugo et al., 2014).

Before diving into contrasts between the effects of *rho* and *rpoB* mutations, it is useful to briefly consider the structure and function of RNAP and Rho. RNAP consists of a five-subunit core protein that has the catalytic activity for RNA synthesis (Ebright, 2000, Ishihama, 2000). The *β* subunit contributes to RNA synthesis activity and contains regions for non-sequence-specific interactions with DNA and nascent RNA. Some of the observed beneficial *rpoB* mutations have occurred in the contact region between the downstream DNA duplex and RNAP (proximal active site) (Rodríguez-Verdugo et al., 2014). Mutations in the contact region are particularly likely to play key roles in modulating RNAP activity in the three stages of transcription: Initiation, elongation, and termination (Ederth et al., 2006).

The Rho molecule acts to terminate RNAP transcription for a subset of *E. coli* genes. The protein is a hexameric ring composed of six monomers (Skordalakes and Berger, 2003) (fig. 1). Given a mature hexamer, transcriptional termination proceeds, first, by the protein recognizing a *rho ut*ilization (*rut*) site on an elongating mRNA; second, by binding *rut* sites and translocating mRNA through the hexamer’s central cavity (Skordalakes and Berger, 2003); until, third, it contacts RNAP while it pauses, thereby disassociating the elongation complex (Skordalakes and Berger, 2003, Peters et al., 2011). The *rut* sites remain difficult to predict bioinformatically, and so the genomic targets of Rho are characterized incompletely (Ciampi, 2006, Hollands et al., 2014). However, a few hundred intergenic and intragenic Rho termination sites have been identified experimentally (Peters et al., 2009, 2012), and these have helped to evaluate whether *rho* mutants increase (Mori et al., 1989, Miwa et al., 1995) or decrease the efficiency of termination (Freddolino et al., 2012, Haft et al., 2014).

**FIG. 1.**
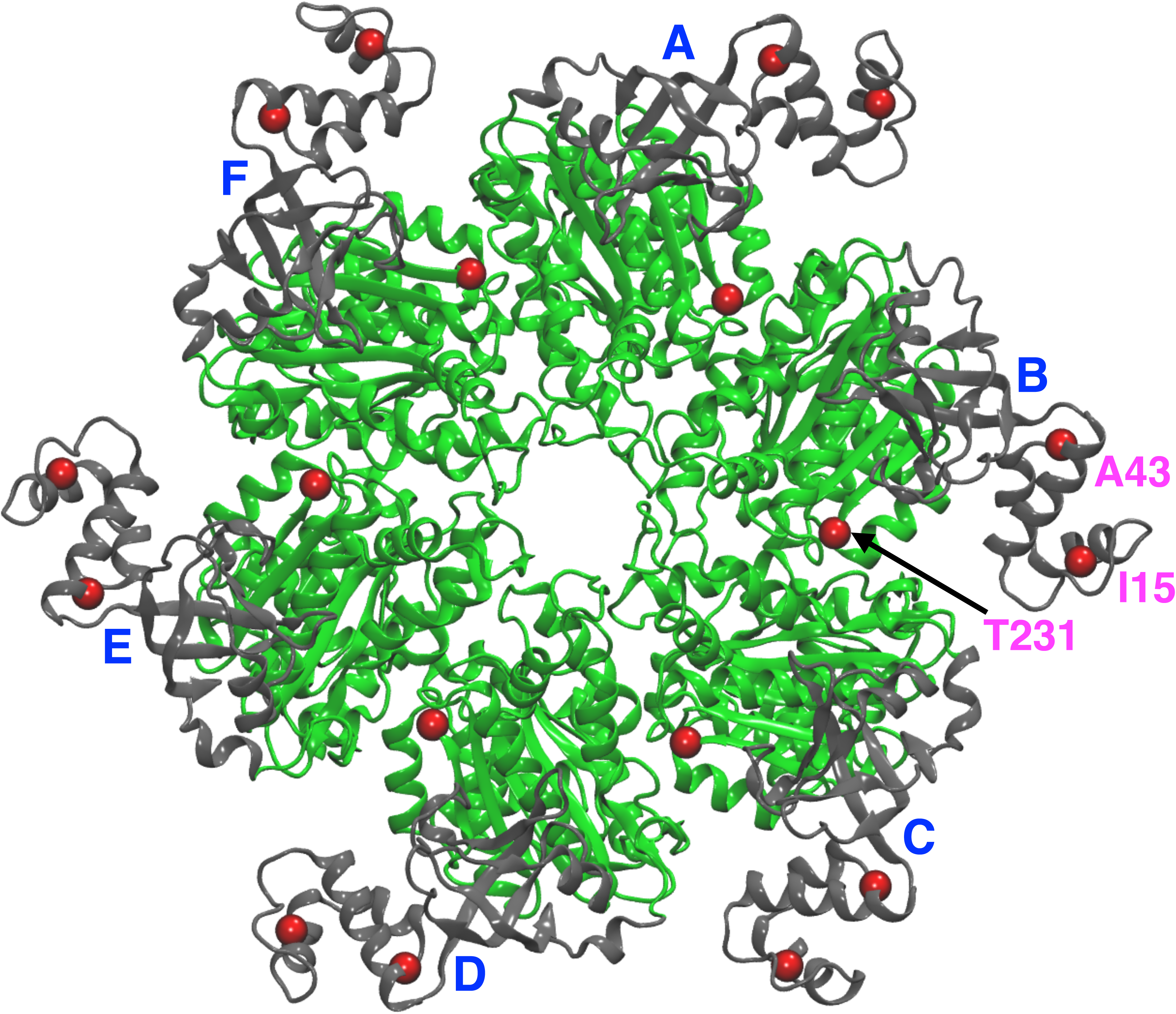
A schematic of the Rho protein, showing each of its six hexamers (A-F) and the location of the single-mutations (red spheres) produced for this experiment. Mutations in *I15* and *A43* are located in the N-terminal domain (NTD) (gray); the mutation in *A231* in the C-terminal domain (CTD) (green).

Here we take advantage of a unique opportunity to compare and contrast the effect of adaptive mutations between the *rho* and *rpoB* genes that characterize alternative adaptive pathways. In the study, we engineer four beneficial *rho* mutations into the REL1206 ancestor and contrast effects on fitness, growth, gene expression and transcriptional termination to four previously-studied *rpoB* mutants (Rodriguez-Verdugo et al., 2014, Rodríguez-Verdugo et al., 2016). In doing so, we address three sets of questions. First, are the fitness and growth characteristics similar between *rho* and *rpoB* mutations? Second, do the beneficial mutations influence the expression of similar numbers and sets of genes? Third, do *rho* mutations alter its termination efficiency and, if so, how do termination properties compare between *rho* and *rpoB* mutations? Finally, do the mutations have a consistent predicted effect on protein stability, which is predicted to be linked to fitness (e.g. DePristo et al., 2005)?

## RESULTS

### Rho mutations and relative fitnesses

We engineered four *rho* mutations - *I15N*, *I15F, A43T* and *T231A* – into the ancestral REL1206 background (see Materials and Methods). Each of these substitutions were adaptive by the criterion of parallelism, because they were observed in 15, 2, 3 and 2 independent populations, respectively, of our 42.2°C thermal stress experiment (Tenaillon et al., 2012). *I15N*, *I15F* and *A43T* point mutations are located in the NHB domain of the Rho monomer, while *T231A* is located in the central part of the C-terminal domain (CTD), specifically in the Loop-CTD (fig. 1).

We assessed the relative fitness 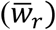 of the four *rho* mutants to REL1206 at 42.2°C (supplementary table S1, Supplementary Material online). Relative fitness data conveyed three observations (table 1). First, two of the four mutations (*A43T* and *T231A*) yielded a 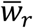value significantly > 1.0, indicating an adaptive benefit under thermal stress. In contrast, neither the *I15N* nor the *I15F* mutation produced a 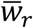value significantly different from 1.0. *I15N* yielded an estimate < 1.0 (Rodriguez-Verdugo et al., 2014) suggesting a slightly deleterious mutation (table 1). Second, the 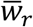estimates differed significantly among the four mutations (ANOVA, *p* = 2.4 × 10^-8^), with *I15N* and *I15F* forming one group and *A43T* and *T231A* forming a second, significantly different group (Tukey-test; *p* < 0.05).

**Table 1.** Relative fitness measured at 42.2°C for four *rho* mutants and four *rpoB* mutants, along with their estimated ΔΔG values, as predicted by molecular modeling.

| Mutant<sup>1</sup> | Relative Fitness | | | Estimated $\Delta\Delta G$ values | |
| --- | --- | --- | --- | --- | --- |
| | Mean | ± 95% CI | *p*-value<sup>2</sup> | Location<sup>3</sup> | $\Delta\Delta G$ at 42.2°C<br>(kcal/mol) ± S.E.<sup>4</sup> |
| *rho I15F* | 1.008 | 0.117 | 0.837 | $\alpha 2$ NHB<sup>*</sup> | 1.22 ± 0.25 |
| *rho I15N* | 0.945 | 0.184 | 0.396 | $\alpha 2$ NHB<sup>*</sup> | 0.50 ± 0.35 |
| *rho A43T* | 1.237 | 0.093 | < 0.05 | $\alpha 3$ NHB<sup>*</sup> | 1.17 ± 0.39 |
| *rho T231A* | 1.266 | 0.104 | < 0.05 | Loop - CTD<sup>**</sup> | -0.06 ± 0.45 |
| *rpoB I572F* | 1.155 | 0.047 | < 0.05 | Rif<sup>+</sup> | -0.327 ± 0.59 |
| *rpoB I572L* | 1.229 | 0.099 | < 0.05 | Rif<sup>+</sup> | -0.421 ± 0.26 |
| *rpoB I572N* | 1.184 | 0.067 | < 0.05 | Rif<sup>+</sup> | 0.803 ± 0.52 |
| *rpoB I966S* | 1.258 | 0.106 | < 0.05 | Eco flap - βi9<sup>++</sup> | 1.405 ± 0.34 |

<sup>1</sup> *rho* data from this study, based on nine replicated measures. *rpoB* data include that from (Rodriguez-Verdugo et al., 2014) plus three additional replicates, for a total of six replicates
<sup>2</sup> $p < 0.05$ rejects the null hypothesis that $\bar{w}_r = 1.0$ , indicating that the mutation is beneficial ( $\bar{w}_r > 1.0$ ) relative to the REL1206 ancestor.
<sup>3</sup> Location of mutation in the Rho protein: NH B=N-terminal helix bundle \* Mutations located in the NTD= N-terminal and \*\*CTD= C-terminal domain. Location of mutations in the RNAP protein: <sup>+</sup> RNA polymerase, $\beta$ subunit, Rif= Rif-binding regions in the fork domain, <sup>++</sup> RNA polymerase, $\beta$ subunit, Eco flap domain, $\beta$ subunit insert 9.
<sup>4</sup> Errors are calculated with 95% confidence interval. $\Delta\Delta G$ value for *rho* and *rpoB* W.T. is zero.

We contrasted 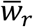 among four *rpoB* mutants (*I572L*, I*572F*, *I572N*, *I966S*) (Rodriguez-Verdugo et al., 2014) and the four *rho* mutants. Each of the *rpoB* mutations had 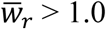, with estimates ranging from 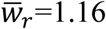 to 1.26 at 42.2°C (table 1). Across all mutants, comparisons again indicated heterogeneity of 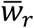 (ANOVA, *p* = 1.91 × 10^-08^; table 1), and Tukey tests grouped *I15N* and *I15F* against the remaining six mutations (*p* < 0.05 for 10 of 12 comparisons between *I15* mutations and the remaining mutations). Altogether, 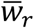 values were >1.00 and indistinguishable among the four *rpoB* mutations and two of the four *rho* mutations; the two remaining *rho* mutations had demonstrably lower 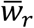 values and little effect on fitness.

### Growth dynamics

To gain additional insights about potential differences between *rho* and *rpoB* mutants, we measured growth curves of mutant lines at 37.0°C and 42.2°C (fig. 2, table 2 and supplementary table S2, Supplementary Material online) and estimated maximal growth rates and final yields at stationery phase. At 37.0°C, all *rho* mutants had significantly higher final yields than at 42.2°C (supplementary table S2, Supplementary Material online), indicating that high temperature remains stressful even after the introduction of putatively beneficial mutations. At 42.2°C, the *rho* mutants had a lower final yield than the ancestor, and there were also differences in maximum growth rate among mutants: *I15F* and *I15N* showed somewhat lower maximum growth rate than the ancestor but *A43T* and *T231A* had higher growth rates (fig. 2 and supplementary table S2, Supplementary Material online), consistent with their higher 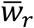 values.

**Table 2.**
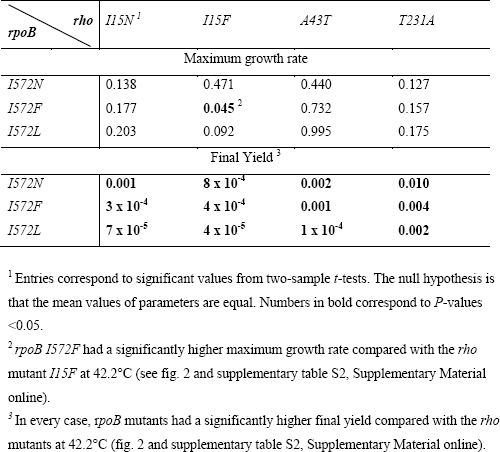
*P*-values for statistical comparisons of maximum growth rate and yield of *rho* and *rpoB* mutants at 42.2°C

**FIG. 2.**
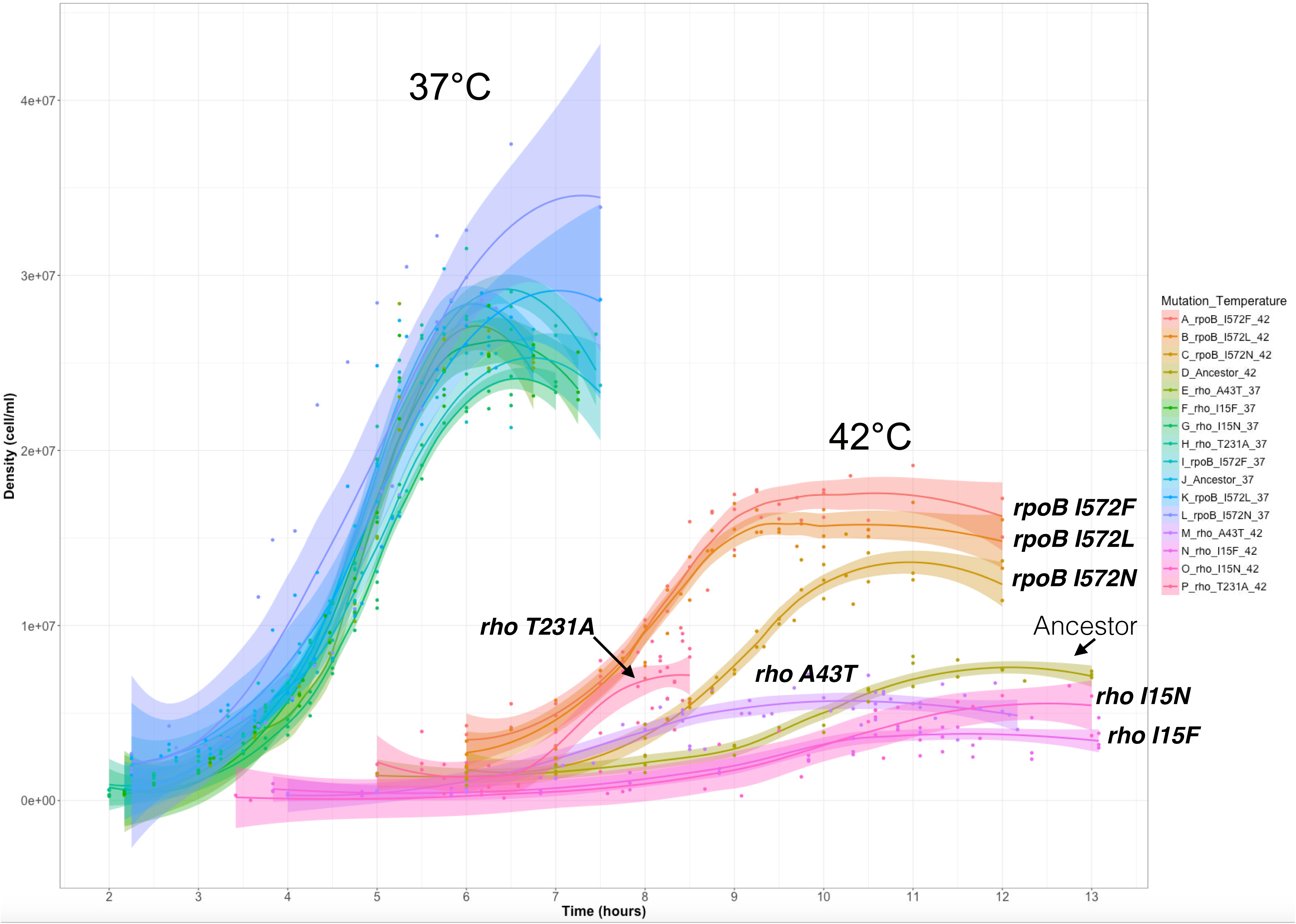
Growth curves for the mutants and ancestor at 37.0°C and 42.2°C. Each line corresponds to a local polynomial regression fitting of three replicates with its 95% confidence interval (see table 2 and supplementary table S2).

We then contrasted growth dynamics between *rho* and the three *I572 rpoB* mutants (Rodríguez-Verdugo et al., 2016). Similar to results with *rho*, *rpoB* mutants grew more slowly and had lower yields at 42.2°C compared to 37.0°C (fig. 2 and supplementary table S2, Supplementary Material online). At 37.0°C, both *rho* and *rpoB* mutants grew with similar rates relative to the REL1206 ancestor (fig. 2 and supplementary table S2, Supplementary Material online). However, a striking difference became apparent at 42.2°C, because the *I572 rpoB* mutants reached significantly higher final yields than the *rho* mutants (table 2 and supplementary table S2, Supplementary Material online). The only significant difference in growth rates was the contrast between *rpoB I572F* vs. *rho I15F* contrast (table 2). Overall, growth curves indicate that: *i*) *rho* and *rpoB* mutations differ with respect to yield, and *ii*) growth rates tend to be lower for *rho I15F* compared to the remaining clones.

### Gene expression phenotypes

To gain further insight into the phenotypic effects of *rho* mutations, we measured gene expression at 42.2°C using replicated RNAseq data (see Materials and Methods). We hypothesized that expression covaries with 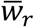, so that *I15N* and *I15F* differ more dramatically in expression from *A43T* and *T231A* than they do from each other. To test this hypothesis, we measured the correlation in expression across all genes for each pair of *rho* mutants (table 3). All pairwise correlations were high (*r*^*2*^ *>* 0.95), but *A43T* and *T231A* had the highest correlation (*r*^*2*^ = 0.992), while contrasts between an *I15* mutant with either *A43T* or *T231A* had lower correlations (table 3). We also identified differentially expressed genes (DEGs) between *rho* mutants, based on a significance cut-off of *q* < 0.001 (see Material and Methods). Only one gene was expressed differentially between *I15F* and *I15N*, but both *I15* mutations had > 100 DEGs between *A43T* and *T231A* (table 3). These data suggest that *A43T* and *T23A* mutations have quantitatively different effects on GE relative to the two *I15* mutations, mimicking observed differences in fitness among clones.

**Table 3.** Comparisons of gene expression between single *rho* mutants. The cells above the diagonal report the Pearson correlation coefficient in gene expression across all genic regions (*n*=4204). The cells below the diagonal indicate the number of genes significantly different via DEseq (*q* <0.001).

|  | *I15N* | *I15F* | *A43T* | *T231A* |
| --- | --- | --- | --- | --- |
| *I15N* |  | 0.991 | 0.962 | 0.955 |
| *I15F* | 1 |  | 0.969 | 0.964 |
| *A43T* | 149 | 190 |  | 0.992 |
| *T231A* | 419 | 552 | 16 |  |

Given that single *rho* mutations can confer extensive GE shifts relative to an ancestral background (Freddolino et al., 2012, Haft et al., 2014), we compared GE between the REL1206 ancestor at 42.2°C and each of the four *rho* mutants. Across the four *rho* mutants, we detected a total of 1140 DEGs compared to the 42.2°C ancestor, but there were more DEGs for *T231A* (1028 genes) and *A43T* (656) than for *I15F* (372) and *I15N* (195) (fig. 3*A*). Among the complete set of 1140 DEGs, 141 (or 12%) were shared among all four *rho* mutants (fig. 3*A*). Gene Ontology (GO) analysis of these 141 genes revealed significant enrichment for down-regulation of maltose transport genes (GO: 0015768, *p* = 1.04 x10^-3^) and significant up regulation of genes involved in transcription (GO:0010467, *p* = 1.12 × 10^-12^) and translation (GO: 0006412, *p* = 9.17 × 10^-37^) (supplementary table S3, Supplementary Material online). Interestingly, the *A43T* and *T231A* mutants shared 330 DEGs that were not identified in either *I15F* or *I15N*. These 330 genes are likely candidates to contribute to 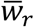 differences among *rho* mutants, and they were enriched for up-regulation of purine nucleotide biosynthetic processes (GO: 0006164, *p* = 1.15 × 10^-12^) and down-regulation for glycerol catabolic processes (GO: 0019563, *p* = 1.15 × 10^-12^) (supplementary table S4, Supplementary Material online).

**FIG. 3.**
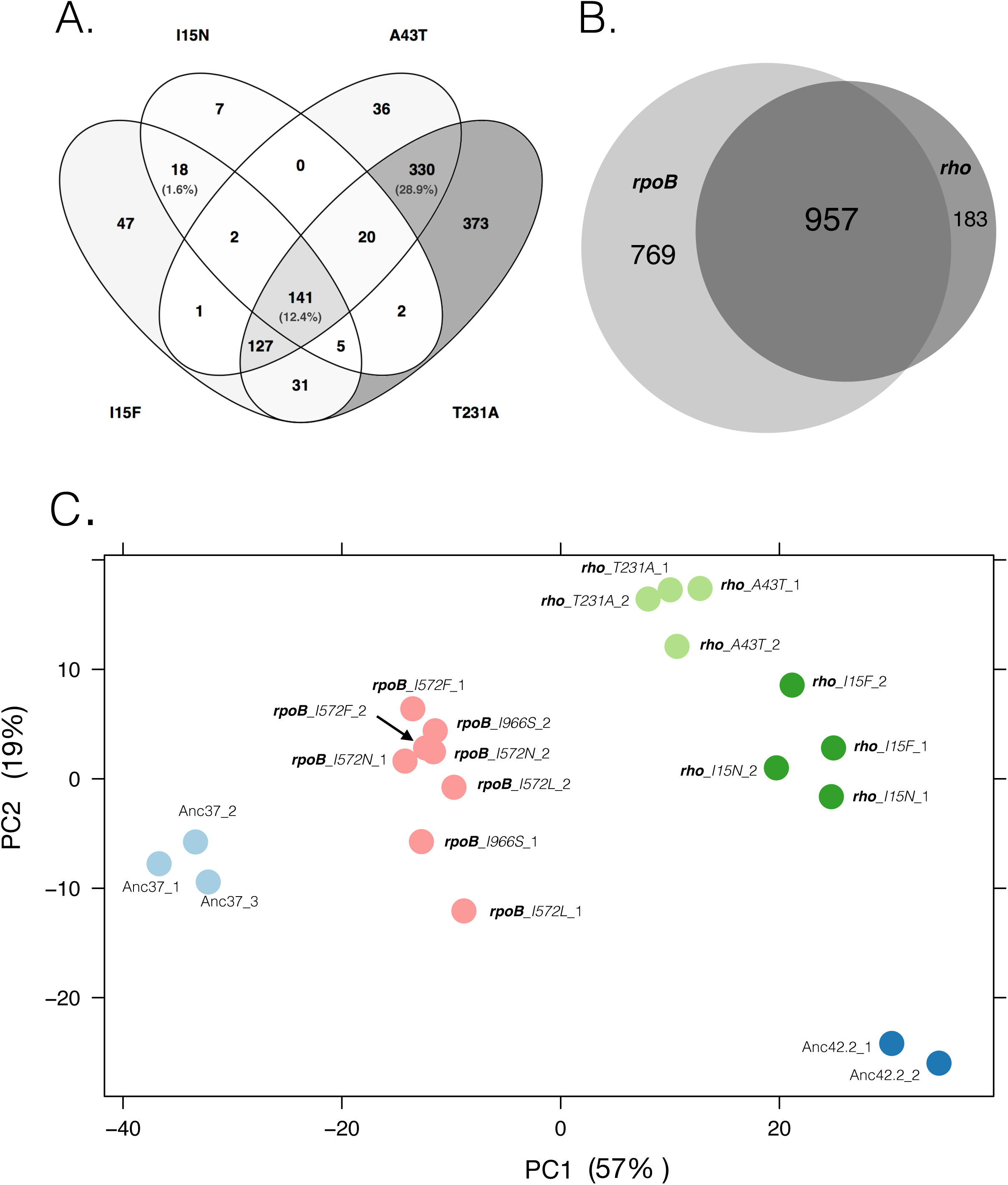
(*A*) A Venn diagram of the number of genes that exhibit significant DEG between *rho* mutants. *(B)* A Venn diagram of the number of genes that differ in expression between the four *rho* mutants and REL1206 at 42.2°C and all four *rpoB* mutants and REL1206 at 42.2°C. *(C)* A PCA plot of the RNAseq data, including replicated data from REL1206 at 37.0°C (light blue dots), REL1206 at 42.2°C (dark blue dots); replicated data from each of the four *rho* mutations (light and dark green dots); and replicated data from each of the four *rpoB* mutations (pink dots). The *x*-axis represents restoration along PC1, where smaller values reflect greater restorative effects.

Finally, to determine whether putatively beneficial mutations in *rpoB* or *rho* have similar effects on gene expression, we identified DEGs in four *rpoB* mutants -- *I572N*, *I572F*, *I572L* and *I966S* -- using previously published data (Rodríguez-Verdugo et al., 2016). We took the union of the DEGs for the four *rpoB* mutations and found 1726 DEGs relative to REL1206 at 42.2°C. The *rpoB* and *rho* DEG sets were remarkably overlapping (fig. 3*B*): of the 1726 DEGs identified with *rpoB* mutations, 957 genes (or 55.4%) were also identified as DEGs in *rho* mutants. This similarity is more impressive when one considers that *rho* mutations altered gene expression for significantly fewer genes than *rpoB* mutations (1140 vs. 1726; binomial test: *p* < 2.2 × 10^-16^) so that 83% (=957/1140) of DEGs caused by *rho* mutations were also identified as DEGs in *rpoB* mutants (fig. 3*B*). These results indicate that *rho* mutants generally affect the expression of a subset of the genes affected by *rpoB* mutants.

The DEG overlap between *rho* and *rpoB* mutations left only 183 DEGs unique to the set of *rho* mutations (fig. 3*B*). Among these 183 DEGs, GO analyses identified up-regulated responses to temperature stimuli (e.g., *clpB*, *dnaK*, *dnaJ*, *degP, grpE*; GO: 0009408, *p* = 3.35 x10^-2^) (supplementary data set S1, Supplementary Material online). In contrast, the 769 genes uniquely modified by *rpoB* mutants were enriched for up-regulation of iron ion transport (GO: 0006826, *p* = 3.2 × 10^-05^) and flagellum cell motility (GO: 00048870, *p* = 7.1 × 10^-06^) genes. Monosaccharide catabolic processes (GO: 0005996, *p* = 2.0 × 10^-04^) and cell wall biogenesis (GO: 0042546, *p* = 2.94 x10^-4^) were also unique to *rpoB* mutants, but they were not consistently up- or down-regulated (supplementary data set S1, Supplementary Material online).

### Shifts in gene expression are partially restorative

Taken in total, *rpoB* and *rho* mutants affect an overlapping gene set, but it is not a foregone conclusion that they affect expression in the same direction. Here direction refers to expression shifts in mutants relative to the stressed ancestral state (i.e., REL1206 at 42.2 °C) and the unstressed ancestral state (i.e., REL1206 at 37.0°C) (Carroll and Marx, 2013). We determined directionality in two ways. First, we applied PCA to all expression data for *rho, rpoB* and ancestral replicates, and then plotted the first two principal components, PC1 and PC2. Based on the PCA, the *rpoB* mutants were intermediate on PC1 between the ancestor at 37.0°C and 42.2 °C (fig. 3*C*). This position is consistent with an overall tendency of *rpoB* mutants to restore expression patterns from the stressed (42.2°C) toward the unstressed (37.0°C) ancestral state (Rodríguez-Verdugo et al., 2016). Expression patterns from *rho* mutants were also intermediate between the two ancestral states on PC1, but they were not restored as fully as the *rpoB* mutants (fig. 3*C*). Intriguingly, replicates from the two less fit *rho* mutants (*I15N* and *I15F*) clustered more closely to REL1206 at 42.2°C. These clustering patterns suggest that fitness is related to restoration in PCA space; indeed, 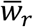and the average PC1 eigenvector were correlated across the eight mutants (*r*^*2*^ = 0.4156; *p* = 0.042).

Second, we categorized expression of single genes into categories. Following previous studies (Carroll and Marx, 2013), genes in mutants were classified as: *i*) “restored”, if expression in the mutant shifts toward the unstressed ancestral state from the stressed ancestral state; *ii*) “unrestored”, if expression remains similar to the REL1206 ancestor in its stressed state; *iii*) “reinforced” if expression in the mutant is exaggerated in the mutant relative to REL1206 at both 37.0°C and 42.2°C and, finally, *iv*) “novel” if the two ancestral treatments did not differ in GE but the mutant differed from both. (The Materials and Methods section includes more precise quantitative definitions of these four directional categories.) The majority of genes fell into the restored category for all mutants except *rpoB I572L* (table 4). More importantly, the number of restored genes was correlated with 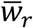 across the eight mutants (*rho* and *rpoB* mutants) at a borderline level of significance (Spearman’s *r* = 0.67; *p* = 0.055). Altogether, directional analyses show that 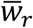of a mutant is a function of the degree of restoration of gene expression.

**Table 4.** The direction of gene expression for genes within *rho* and *rpoB* mutants.

| Mutant <sup>1</sup> | Restored <sup>2</sup> | Unrestored | Reinforced | Novel | Total <sup>3</sup> |
| --- | --- | --- | --- | --- | --- |
| *rho I15F* | 762 | 886 | 19 | 4 | 1671 |
| *rho I15N* | 802 | 857 | 8 | 16 | 1683 |
| *rho A43T* | 1115 | 544 | 7 | 32 | 1698 |
| *rho T231A* | 1250 | 405 | 12 | 46 | 1713 |
| *rpoB I572F* | 1092 | 644 | 1 | 59 | 1796 |
| *rpoB I572L* | 563 | 1174 | 0 | 6 | 1743 |
| *rpoB I572N* | 1165 | 569 | 3 | 53 | 1790 |
| *rpoB I966S* | 1308 | 428 | 1 | 39 | 1776 |

<sup>1</sup> *rho* data from this study. *rpoB* data from (Rodriguez-Verdugo et al., 2014).
<sup>2</sup> The number of genes was counted as restored, unrestored, reinforced and novel using the rules elaborated in the Material and Methods.
<sup>3</sup> The total tallies the number of genes that differ at $q < 0.001$ from either REL1206 at 37.0°C or REL1206 at 42.2°C or both. The total differs significantly between *rho* and *rpoB* mutations (t-test; $p = 0.0016$ ).

### Assessing potential effects on transcriptional termination

We have shown that *rho* mutations differ in 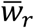 and that 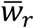 correlates with shifts in the magnitude and direction of gene expression. However, the question of mechanism remains – i.e. how do mutations in *rho* affect its function as a transcriptional terminator? Previous studies have shown that *rho* mutations tend to decrease termination and therefor increase read-through activity (Freddolino et al., 2012, Haft et al., 2014). Alternatively, we have hypothesized that adaptive *rho* mutations increase transcriptional efficiency at high temperature by enhancing termination (Rodriguez-Verdugo et al., 2014, Rodríguez-Verdugo et al., 2016). We addressed this hypothesis both directly and indirectly.

#### Direct effects on Rho-terminated regions

The direct approach relied on a set of 178 Rho-terminated regions that were defined by Peters et al. (2009) and previously used to analyze read-through activity of a *rho* mutation (Haft et al., 2014) (supplementary data set S2, Supplementary Material online). The advantage of studying these regions is that they can, in theory, examine direct (primary) effects of *rho* mutations, as opposed to indirect (secondary) effects on downstream genes. The size of the *rho*-terminated regions spanned from 100 to ≈ 4000 bp, with roughly half located at the 3’ end of coding regions (intergenic) and the other half within genes (intragenic). Following Haft et al. (2014), we calculated *γ*, the log_2_ ratio of normalized RNAseq counts between each mutant at 42.2°C and REL1206 at 42.2°C for each of the 178 regions (see Materials and Methods). Under the null hypothesis that the mutant and ancestor have similar termination properties, *γ* is distributed binomially. If a mutation enhances termination, we expected significantly more than 50% of the regions to have *γ* < 0.0. In contrast, increased read-through results in >50% of regions having *γ* > 0.0 (Haft et al., 2014). Contrary to our prediction, both *I15N* and *I15F* have increased read-through activity, with 108 and 119 of 178 regions having *γ* > 0.0 (binomial, *p* < 0.01). The two remaining *rho* mutations (*A43T, T231A*) yielded no evidence of altered termination properties (*p* = 1.00; supplementary fig. S1, Supplementary Material online)

We further examined expression dynamics of the Rho-terminated regions by taking the ratio of expression counts for each *rho*-terminated region (RT) against the counts from the immediate 5’ upstream region (UP) of the same size. We then computed a measure of termination efficiency for each mutant as the ratio of averages, *r*_*avg*_, of read counts in RT vs. UP across all 178 regions (see Materials and Methods). While not without caveats (such as the fact that upstream and downstream regions are in different operons in some cases), this approach allowed us to measure the strength of termination for each mutant and for each ancestral treatment.

Bootstrapped estimates of *r*_*avg*_ are presented in fig. 4. The estimates suggest weaker termination transcription (i.e., enhanced read-through) for REL1206 at 42.2°C compared to 37.0°C; although the trend is notable, the two ancestral treatments do not differ significantly (bootstrap; *p* =0.864). For the *rho* mutants, *I15N* was alone in having a higher median estimate of *r*_*avg*_ than REL1206 at 42.2°C (fig. 4), which verifies the inference of increased read-through activity based on *γ* values. In contrast, the *r*_*avg*_ estimates for *T231A* and *A43T* (and *I15F*, to a lesser extent) appeared to restore termination to the level of the ancestor at 37.0°C (fig. 4).

**FIG. 4.**
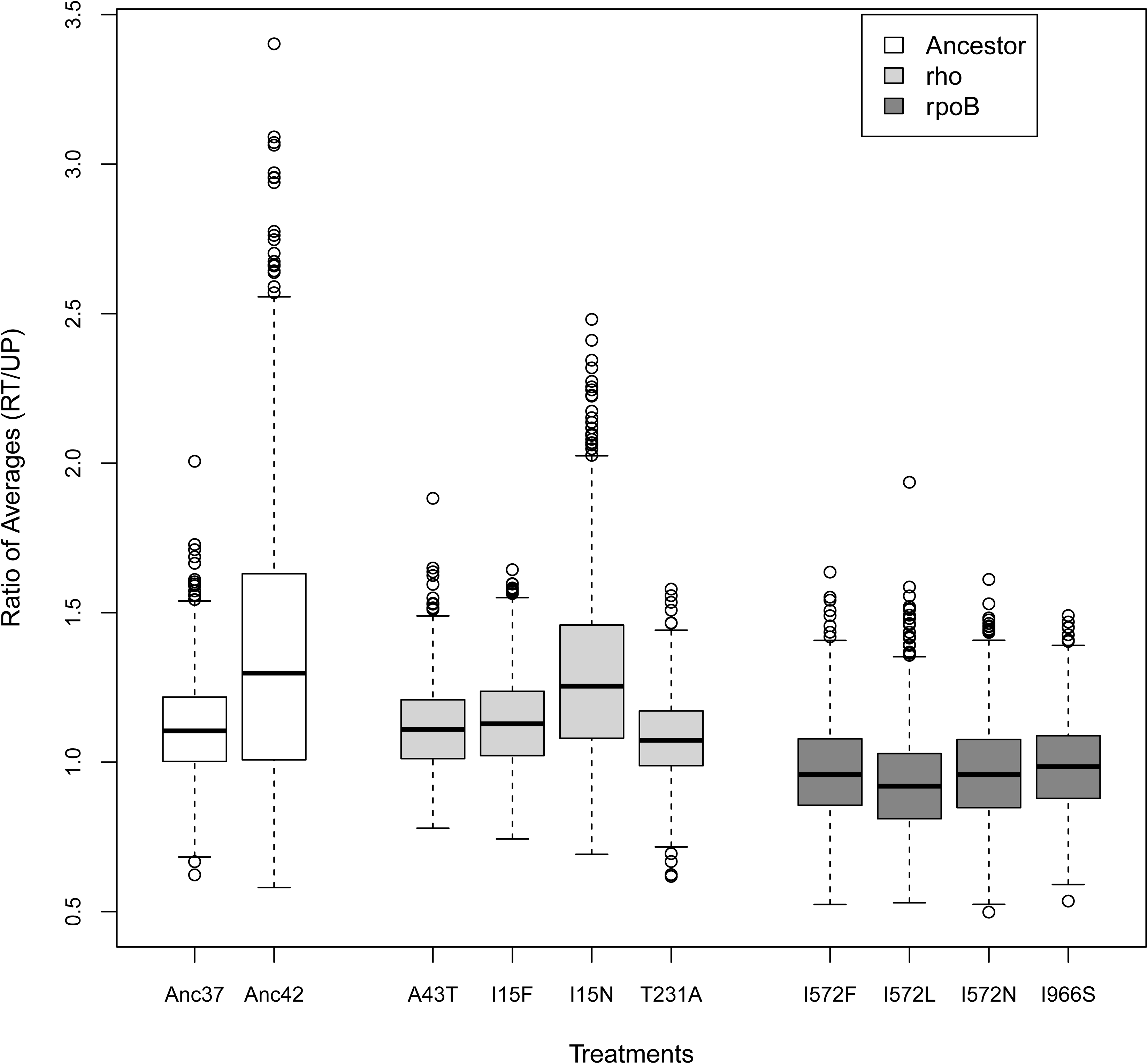
A graph of bootstrap estimates of the ratio of averages 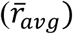 that measures the ratio of counts between *rho* terminated sites (RT) and their upstream 5’ regions (UP) across 178 loci. Lines in the boxplots represent the median bootstrap-based estimate of 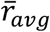 based on 1,000 resamplings (see Material and Methods). The boxes represent the upper and lower quartiles of bootstrap replicates. Anc37 represents the REL1206 ancestor at 37.0°C and Anc42 is the ancestor at 42.2°C.

It has been shown previously that some rpoB mutants enhance transcriptional termination (Jin et al. 1988). Hence for completeness we also applied analysis of Rho-terminated regions to *rpoB* mutants. Based on *γ* analyses, none of the *rpoB* mutants yielded evidence of altered termination in these regions (*p* > 0.500; supplementary fig. S1, Supplementary Material online). However, *r*_*avg*_ estimates for *rpoB* were consistently lower than the ancestral treatments and the *rho* mutants (fig. 4). Furthermore, the average *r*_*avg*_ estimate between the set of *rho* and *rpoB* mutants differed significantly (*p* < 0.001), indicating either that termination in these regions is more efficient in *rpoB* mutants or that *rpoB* mutants had consistently higher expression in the upstream regions. The *rpoB* mutations also differed significantly from the ancestors at 42.2°C and 37.0°C (*p*<0.003). Altogether, *r*_*avg*_ and *γ* analyses indicate that the *rho I15N* and *I15F* mutations likely decrease termination efficiency, but as a group the *rpoB* mutations have statistically better termination for these same regions.

#### Indirect effects

In addition to assessing the properties of *rho* termination directly, we assessed potential effects on termination indirectly by focusing on the expression of all genes and intergenic regions (IRs). Focusing first on genes, we compared expression data between REL1206 at 37.0°C and 42.2°C. Similar to Rodriguez-Verdugo et al. (2016), we found that ∼1700 of 4204 coding regions were differentially expressed (*q* < 0.001) during thermal stress, with a tendency toward lower expression at 42.2 °C (972 down-regulated genes vs. 695 up regulated genes; binomial *p* = 1.243 × 10^-11^) (fig. 5*A*). Given this baseline, we then assessed the effect of *rho* mutants relative to REL1206 at 42.2°C. GE was significantly biased toward higher expression in each of the mutants relative to REL1206 at 42.2°C at *q* < 0.001 (fig. 5*A*; binomial, *p* = 1.41 × 10^-10^ (*I15N*), *p* = 2.13 × 10^-6^ (*I15F*), *p* = 8.6 × 10^-12^ (*A43T*), *p* = 2.2 × 10^-16^ (*T231A*)). In effect, thermal stress led to lower GE at 42.2°C relative to 37.0°C, and the four mutations reversed that trend. These trends were quantitatively similar for *rpoB* mutants (fig. 5*C*)

**FIG. 5.**
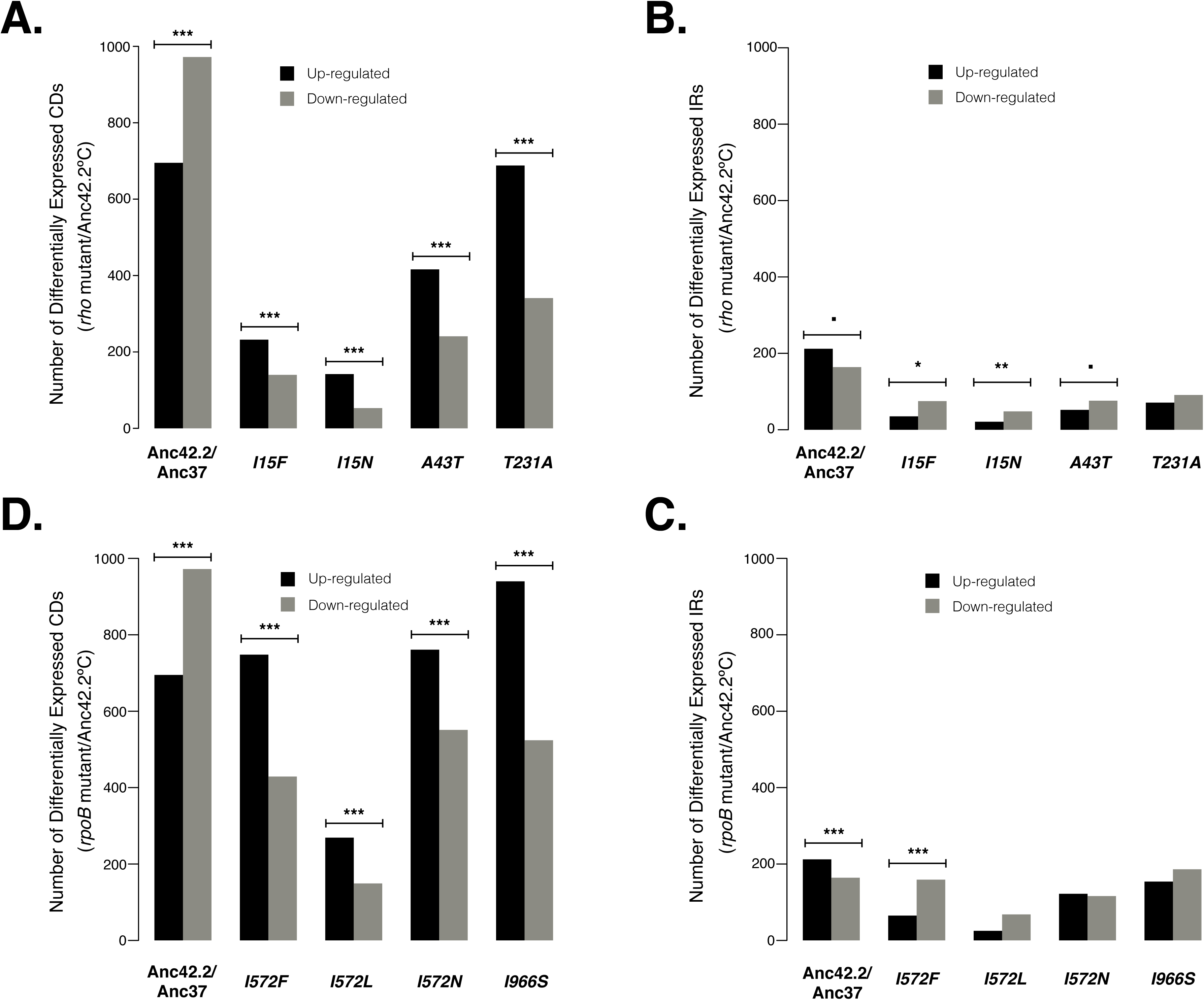
Genome-wide expression patterns of genic and intergenic regions at 42.2°C. Histograms showing the number of coding regions that are up or down regulated compared to REL1206. *(A)* The first pair of histograms shows REL1206 at 37.0°C vs. 42.2°C. Up- or down-regulation is relative to the first of the listed pair. For example, the first pair of histograms show that REL1206 at 42.2°C has ∼700 and ∼1000 genes that are expressed at higher and lower levels, respectively, than REL1206 at 37.0°C. The remaining histograms compare *rho* mutants at 42.2°C to REL1206 at 42.2°C. The bars with three, two and one asterisks indicate *p* < 0.0001, *p* < 0.001 and *p* < 0.01 respectively, a dot indicates *p* < 0.05. *(B)* Similar histograms as in *A*, but based on intergenic regions (IRs). *(C)* and *(D)* are identical to *(A)* and *(B)* but based on the *rpoB* mutants.

The picture was, however, different for IRs. Among 2306 IRs (supplementary data set S3, Supplementary Material online) we found more were up-regulated in REL1206 at 42.2°C relative to 37.0°C (212 up regulated IR vs. 164 down; regulated; binomial *p* = 0.015) (fig. 5*B*). This up-tick in the expression of IRs may indicate more transcriptional read-through for REL1206 under thermal stress. This trend was reversed by both *rho* and *rpoB* mutations, such that differentially expressed IRs tended to have *lower* expression in the mutants relative to REL1206 at 42.2°C [fig. 5*B*, binomial *p* = 0.0016 (*I15N*), *p* = 0.0001 (*I15F*), *p* = 0.04164 (*A43T*); fig 5*D*; *p* = 0.0016 (*I15N*), *p* = 0.0001 (*I15F*), *p* = 0.04164 (*A43T*)]. These results raise the possibility that all of the *rho* and *rpoB* mutations enhance termination efficiency at 42.2°C. For the *rho* mutations this is not likely a direct effect (fig. 4) but an indirect, downstream effect.

### Mutations and protein stability

Finally, we evaluated the predicted folding stability of the Rho and RpoB proteins. Our principal goal was to assess whether changes in fitness correlated with the direction and magnitude of shifts in the free energy of protein folding stability, as is expected under some evolutionary models (DePristo et al., 2005, Wylie and Shakhnovich, 2011). We hypothesized that 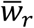 from specific mutations correlates with the difference of the free energy of folding between ancestral and mutant proteins (ΔΔG) at 42.2^o^ C. To determine the free energy of folding (ΔG), we applied an approach that combines classical Molecular Dynamics (MD) with the FoldX algorithm (see Materials and Methods) and then measured the change in folding stability (ΔΔG) induced by each of the four *rho* and four *rpoB* mutations. Positive values of ΔΔG suggest a decrease either in the stability of the protein fold or in the ability of the protein to fold relative to the ancestor; negative values indicate increased stability (Guerois et al., 2002, Miller et al., 2016). Three of four *rho* mutations (*I15F*, *A43T* and *I15N*) were destabilizing at 42.2°C, with ΔΔG values ranging between 0.50 to 1.22 kcal/mol (table 1); in contrast, the *T231A* substitution mildly increased folding stability at 42.2°C (ΔΔG=-0.06). Two of the four *rpoB* mutations were destabilizing at 42.2°C (table 1). Although our sample size was small (*n* = 4) for each gene, there was no clear trend between 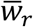 and ΔΔG (table 1). For completeness, we estimated ΔΔG for 16 additional *rho* mutations from our evolution experiment, for a total of 20 ΔΔG estimates (supplementary table S5, Supplementary Material online). Of these, 17 of 20 ΔΔG estimates were destabilizing, and there were no differences in ΔΔG for mutations in different structural features (e.g., α-helices vs. *β*-sheets, t-test, *p* = 0.45; NHB and CSD-like RBD, vs. non-defined regions, *p* = 0.36, see supplementary table S5, Supplementary Material online). The only consistent effect of mutations was toward destabilizing the wild type Rho protein at high temperature.

## DISCUSSION

This study was designed to compare fitness and phenotype, as measured primarily by gene expression, among and between adaptive mutations in *rho* and *rpoB*. We undertook this study both because transcriptional regulators contribute substantially to adaptive evolutionary change across many types of organisms and because mutations of these two genes are particularly common in *E. coli* experimental evolution. However, to our knowledge there have not yet been direct comparisons of the effect of beneficial mutations within and between transcriptional regulators. Our comparisons reveal substantial differences in relative fitness 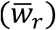 and growth characteristics among mutations, but we also find broad similarities. Specifically, *rho* mutations primarily affect the expression of a subset of the genes affected by *rpoB* mutations, mutations in both genes tend to restore gene and IR expression to the unstressed state of the REL1206 ancestor, and the extent of restoration correlates with 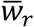.

### Heterogeneity in fitness and growth characteristics

The first major difference among beneficial mutations is in 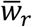. Of the eight mutations, six have 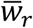 significantly > 1.00 and are statistically indistinguishable (table 1). Of the six advantageous mutations, the three *rpoB* codon *572* mutations are unambiguous drivers of adaptive change, both because they have high fitness values and because they often fix rapidly in experimental populations (Rodriguez-Verdugo et al., 2013).

The two *rho I15* mutations have 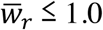 and thus confer no apparent advantage under thermal stress (table 1). These two mutations are unlikely to drive adaptation as solo mutations. They thus illustrate the potential importance of linkage and epistasis, because *I15N* was one of the two most common nucleotide substitutions in the entire thermal stress experiment and therefore must confer a substantial benefit in the proper genetic background. But what is the proper background? Two possibilities stand out: mutations in the *cls* gene or in the *iclR* gene. Knock-downs or knock-outs of the *cls* gene, which encodes a cardiolipin synthase, are present in 14 of 17 of the clones with *rho I15F* and *I15N* mutations; mutations within *iclR*, a transcriptional repressor, are in 10 of 17 of clones; and either *cls* or *iclR* was mutated in all 17 clones (Tenaillon et al., 2012). *rho* mutations have been shown to interact epistatically with mutations in other genes -- such as *rpsL*, *metJ* and *rpsQ* -- that have a role in transcription or translation (Freddolino et al., 2012, Haft et al., 2014). Both *rpsL* and *rpsQ* are critical to the 30S ribosomal unit (Cabezon et al., 1975), and *metJ* represses the expression of genes involved in methionine biosynthesis (Marincs et al., 2006).

The association between *rho* and *iclR* mutations are particularly interesting, because *iclR* encodes a transcriptional repressor of the *aceBAK* operon, which encodes the enzymes involved in the glyoxylate bypass (Nègre et al., 1992). Mutations in *iclR* commonly arise in evolution experiments with *E. coli* that use glucose-limited medium (Herron and Doebeli, 2013; Deatherage et al., 2017), perhaps because they derepress the acetate operon improving the utilization of acetate after glucose is depleted (Spencer et al., 2007). Interestingly, the *aceBAK* operon is up-regulated under thermal stress, but down-regulated again in all of our *rho* and *rpoB* mutants (and significantly down-regulated in *rho A43T*, *T231A* and *rpoB I152F*, *I572N* and *I966S*). Hence, mutations to *iclR* may compensate for the effect of mutations on the *aceB*AK operon (Yamamoto and Ishihama, 2003). It is worth noting that though *iclR* mutations are statistically more likely to be found in the *rho* evolved lines from our thermal stress experiment, they are also found in *rpoB* lines with the *I966*S and *I572N* mutations (Tenaillon et al., 2012). *iclR* mutations may thus compensating for pleiotropic effects.

In contrast to the two *rho I15* mutations, the *A43T* and *T231A* mutations are capable of driving adaptive change as solo mutations, but there is a twist: they too, are often found in *cls* and *iclR* backgrounds. All three *A43T* clones and one of two *T231A* clones from our evolution experiment also had *cls* and *iclR* mutations, again implying the possibility of positively epistatic interactions. Potential functional interactions between *cls* and *rho*, if any, are unknown. The *cls* gene has not been identified as *rho*-terminated (Peters et al., 2009) and is not differentially expressed in any one of our four *rho* mutants. One possible - but purely speculative - interaction between *cls* and *rho* is through membrane signaling. Cardiolipin plays a key role in recruiting signaling molecules to the membrane (Romantsov et al., 2007), and *E. coli* can perceive alterations in membrane fluidity and transfer that information to signal-transduction pathways that alter gene expression (Los and Murata, 2004, Los et al., 2013). The potential for, and causes of, interactions between *rho* and *cls* require further study.

The effects of mutations on growth vary, too (fig. 2). Generally speaking, the *rpoB* mutants grow better at 42.2°C than the *rho* mutants, including higher maximum growth rates and yields (table 2 and supplementary table S2, Supplementary Material online). The effects on maximum growth rates are not statistically significant -- probably due in part to high variance in growth characteristics for the *rho* mutants (fig. 2 and supplementary fig. S2, Supplementary Material online) -- but the final yields for *rpoB* mutations are statistically higher than that of *rho* mutants. Of these two parameters, yield is not a component of fitness (Vasi et al., 1994) and maximum growth rate is at best a proxy for fitness (Concepción-Acevedo et al., 2015, Durão et al., 2015, Wiser and Lenski, 2015). Nonetheless, estimated growth rates corroborate 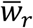 estimates by indicating that the *I15 rho* mutations are not marked improvements (if at all) over the ancestor. Another interesting facet of growth dynamics is that the *rpoB* mutants are robust across 37.0°C and 42.2°C temperature regimes, but the same mutations are antagonistically pleiotropic at lower (< 20.0°C) temperatures (Rodriguez-Verdugo et al., 2014).

### Mutations alter the expression of similar genes in similar directions

Despite differences in relative fitness and growth dynamics between *rho* and *rpoB* mutations, there are some clear similarities with respect to gene expression. One such similarity is that mutations in both genes act in part by restoring expression from the stressed condition (in this case, REL1206 under thermal stress at 42.2°C) back toward the wild type state (in this case, REL1206 at 37.0°C). We have detected this restoration both as a shift in PCA space (fig. 3*C*) and as the dominant directional shift among individual genes (table 4). Several previous studies have also shown that adaptation proceeds through phenotypic restoration (Fong et al., 2005, Carroll and Marx, 2013, Sandberg et al., 2014, Carroll et al., 2015, Hug and Gaut, 2015, Rodríguez-Verdugo et al., 2016), but our study is unique in establishing a quantitative correlation between restoration and 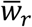.

There is a relevant counter-example. In their study of *Methylobacterium extorquens*, Carroll and Marx (2013) show that adaptation was accompanied by the restoration of ancestral gene expression levels, just as we find here. In their case, growth rates are not correlated with restoration but with reinforcement (i.e., the number of genes that exaggerate the acclimation response to physiological stress). We do not find a correlation between the number of reinforced genes and 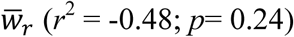. Taken together these studies suggest that relationships among fitness and the direction of evolutionary change may vary with experimental conditions. An important goal for future research is to uncover generalities about relationships among fitness, restoration, reinforcement and novelty.

A second similarity is that the set of genes affected by mutations in *rho* and *rpoB* overlap to a remarkable extent, such that the *rho*-affected genes are a nearly perfect subset of *rpoB*-affected genes (fig. 3*B*). There are, of course, caveats to this observation. One is that to compare the *rho* and *rpoB* pathways in this way, we have taken the union of DEGs among *rho* and *rpoB* mutants. In reality, different mutations are associated with different DEG sets (fig. 3*A*); the pattern is recognizable primarily because we have studied multiple mutants in each gene. A second caveat is to recognize that the number of DEGs is an imperfect metric, because it varies with statistical power and can be non-transitive across pairwise comparisons. Finally, RNAseq approaches are also not without caveats. RNAseq measures relative RNA amounts of expression across genes within a sample, making it difficult to contrast absolute levels of expression between samples. Nonetheless, the overarching impression is that *rho* mutations mostly affect a smaller subset of the genes altered by *rpoB* mutations.

The common set of genes affected by the *rho* and the *rpoB* mutations provides insights into the physiological response to thermal stress. Mutations in *rho* and *rpoB* up-regulate functions related to cellular growth, including purine and pyridine biosynthesis, amino acid biosynthesis, ribosome biogenesis, nitrogen assimilation, translation and the regulation of translation. Mutations in *rho* and *rpoB* also down-regulate a common set of genes, particularly those associated with carbohydrate transport and amino acid degradation. The shared patterns between *rho* and *rpoB* mutations are consistent with the view that long-term acclimation to thermal stress leads to the down-regulation of growth-related pathways, potentially resulting in energy conservation and enhanced survival (Rodríguez-Verdugo et al., 2016) and that adaptation adjusts to support growth in the stressful environment (Gunasekera et al., 2008).

Although there is a considerable overlap of genes affected by *rho* and *rpoB* mutations, some functional categories were altered predominantly by one of the two pathways. For example, *rho* mutants are enriched for DEGs that respond to temperature stimuli, such as *clpB*, *dnaK*, *dnaJ*, *degP* and *grpE.* For reasons not fully understood, the *rpoB* mutations do not up-regulate these temperature response genes (Gunasekera et al., 2008, Rodríguez-Verdugo et al., 2016). In contrast, *rpoB* but not *rho* mutations enhance the expression of iron transport and flagellum biogenesis. This last observation is particularly interesting, because the up-regulation of flagellar genes is unlikely to have been adaptive in our evolution experiment and because two studies have shown that these are later down-regulated by subsequent adaptive mutations in populations (Gunasekera et al., 2008). This example of flagellar genes raises an important point: for all of these gross-scale effects on GE, it is difficult to distinguish between directly adaptive shifts vs. negatively pleiotropic shifts.

### Potential impacts on transcription termination

Genome sequencing uncovers the type and potential function of adaptive mutations in evolution experiments (Long et al., 2015, Tenaillon et al., 2016), but the gap from ‘potential function’ to an established molecular mechanism remains a chasm for all but a few exceptions [e.g., (Blount et al., 2012, Quandt et al., 2015)]. With respect to thermal stress it is known that the speed of RNAP increases with increasing temperature (Ryals et al., 1982). We have therefore hypothesized that adaptive *rpoB* mutations slow RNAP transcription under thermal stress, leading to increased termination efficiency. *rpoB* mutations have been shown to increase termination previously (Jin et al., 1988). The effects of our *rpoB* mutations on the expression of IRs (fig. 5c) and *rho*-terminated regions (fig. 4) are consistent with our increased termination efficiency hypothesis.

Termination efficiency may require a balance (or ‘kinetic coupling’) between Rho and RNAP activity (Jin et al., 1992). Under this model, when RNAP increases in speed, Rho must match the increase to maintain efficient termination. We predicted, then, that adaptive *rho* mutants translocate along RNA more quickly, leading to increased termination efficiency relative to REL1206 at 42.2°C. Improved termination efficiency is possible, in theory, because at least one *rho* mutation (*L3F*) has been shown to increase termination (Mori et al., 1989). Despite this prediction, transcriptomic analyses of *rho* terminated regions indicate that two *rho* mutants (*A43T* and *T231A*) had little detectable direct effect on termination, and the two without a detectable fitness benefit (*I15F* and *I15N*) exhibited weaker termination (i.e., enhanced read-through) (fig. 4).

Our results have at least three implications. First, they imply that a direct shift in Rho termination efficiency – at least as measured here – is not a requirement for adaptation via the *rho* pathway. Second, they unfortunately provide few insights into structure-function relationships, because the free energy of folding (ΔΔG) bears no discernible relationship with read-through properties or with fitness (table 1, supplementary table S5), despite predictions regarding the latter (DePristo et al., 2005, Tokuriki and Tawfik, 2009, Wylie and Shakhnovich, 2011, Bershtein et al., 2012, Serohijos and Shakhnovich, 2014). The only discernible structural pattern was a tendency toward destabilization of the Rho protein (supplementary table S5, Supplementary Material online), which is not surprising given that ∼70% of all possible mutations on globular proteins yield ΔΔG > 1 (Guerois et al., 2002, Schymkowitz et al., 2005) and that beneficial *rpoB* mutations are known to destabilize the *β* subunit of RNAP (Utrilla et al., 2016).

Third, there remains the remarkable result that all of our *rho* and *rpoB* mutations tend torestore gene expression to that of the unstressed ancestor (fig. 2*C*), typically by increasing gene expression (fig. 5*A* and 5*B*) in a similar set of genes (fig. 2*B*). At the same time, all eight mutations tend to reduce expression of IR regions (fig. 5*B* and 5*D*). The latter result is consistent with our hypothesis of enhanced termination efficiency in *rho* mutants. Analyses of *rho*-terminated regions suggest that this is not a *direct* effect of the *rho* mutations but rather an *indirect*, downstream effect. Given the substantial overlap in differentially expressed genes (fig. 2*B*) and intergenic expression patterns (fig. 5*B* and 5*D*), *rpoB* mutations appear to cause a similar effect on transcription termination. For *rpoB* we cannot easily determine whether the effect is direct – i.e., due to a modification of RNAP - or downstream and indirect. Despite our observations, we still do not know how, exactly, enhanced termination efficiency translates to increased fitness and the restoration of gene expression. We can, however, draw several conclusions from our study of *rh*o and *rpoB* mutants: that these beneficial mutations differ in relative fitness, that their fitness is correlated with their effects on gene expression, and that they all, somehow, tend to effect the same sets of genes and simultaneously increase expression of genes while decreasing expression of IR regions. Finally, and perhaps most importantly, these observations show that alternative adaptive pathways encompass substantial phenotypic convergence.

## MATERIALS and METHODS

### The construction of *rho* mutants

The *I15F, A43T and T231A rho* mutations were introduced into the REL1206 ancestral strain using the pJk611 recombineering plasmid, as previously described (Rodriguez-Verdugo et al., 2014). Briefly, the pJk611 plasmid was introduced into REL1206, electroporating 2 µl of plasmid (80 ng of plasmid in total) into 50 µl of competent cells using an Eppendorf Electroporator 2510 set at 1.8 kV. After electroporation, 1 ml LB was added to the electroporated cells, and they were incubated at 30°C for 2 h under constant shaking (120 rpm). Thereafter 100 µl of cells were plated on LB agar plates containing 100 µg/ml ampicillin to select ampicillin-resistant transformants. The ancestral strain carrying the pJk611 plasmid was grown overnight at room temperature (∼20°C) in 25 ml LB with 100 µg/ml ampicillin and 1mM _L-_arabinose (Sigma) until it reached and OD_600_ of 0.6. Electrocompetent cells were made by washing the culture five times with ice-cold water.

To construct each mutant, two oligos of 70 bp were electroporated into cells (supplementary table S6, Supplementary Material online); one oligo introduced the desired single mutation in *rho* and the second oligo incorporated the change that produces an Arabinose positive (Ara+) phenotype. Two µl of each oligo (10 µM) was electroporated into 50 µl of cells. After electroporation 1 ml LB was added, and the cells were incubated at 37.0°C for 3 h with shaking, followed by 500 µl spread onto minimal medium agar (MA) plates supplemented with _L-_arabinose. The plates were incubated 48 h at 42.2°C, after which 188 single colonies were selected and grown on TA agar plates overnight at 37.0°C. We subsequently performed colony PCR, followed by Sanger sequencing to amplify the *rho* gene fragment surrounding the putative mutation (supplementary table S6, Supplementary Material online). PCR conditions were 10 min at 94°C followed by 30 cycles of 20 s at 94°C, 30 s at 60°C, and 1 min at 68°C ending with an extension step of 5 min at 68°C. For each mutation, we isolated multiple lines with the mutation of interest, such that biological replicates were independently derived.

### Relative fitness competitions

We estimated 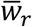by competing mutants against the ancestral REL1206 strain. Briefly, both competitors were revived in LB broth and grown separately overnight at 37.0ºC under constant shaking (120rpm). After cultures were revived, they were subcultured by 1:10000 dilution in 10 ml DM25 media for 24 hours at 37.0°C and 120 rpm and inoculated 1:100 into 9.9 ml of fresh DM25 and incubated at 42.2°C for another day in order to acclimate cells to high temperature. Although the clones had evolved at 42.2°C in DM25, it is common practice in thermal stress studies to allow clones to recover from freezing under less stressful conditions (Bennett and Lenski, 1993, Rodriguez-Verdugo et al., 2014).

Once revived, the two competitors were mixed at a 1:1 volumetric ratio and diluted 100-fold into 10 ml of fresh DM25 and incubated one day at the assay temperature. The initial and final densities were estimated by plating the culture onto tetrazolium-arabinose (TA) agar plates, counting the number of Arabinose+ colonies of the REL1206 ancestor and the number of Arabinose-clones of the mutants and evolved clones. Relative fitness assays were performed on three biological replicates for each mutation and for three technical replicates for each biological replicate; thus each mutation was tested against the ancestor a total of nine times.

ANOVA analyses were based on the *avo* package in R. With ANOVA, we first tested for the effects specific to different biological replicates within a mutation using a model of Fitness∼Mutant*Replicate. Given that there were no differences among biological replicates, the model was simplified to Fitness∼Mutant.

### Growth curves

To obtain the growth parameters of *rho* mutants, we first revived them in LB broth and incubated them overnight at 37.0°C. The overnight cultures were then diluted 10,000 fold into 10 ml DM25 and incubated for 1 day at 37°C. The following day, the acclimated cultures were dilute 100-fold into 10 ml DM25 and incubated for 1 day at the assay temperature (either 37.0°C or 42.2°C). Cell densities were measured every hour during the lag phase and every ∼15 minutes during exponential phase. Cell densities were quantified using an electronic particle counter (Coulter Counter model Multisizer 3 equipped with a 30 µm diameter aperture tube). For this, 50 μl of the growing culture was diluted in 9.9 ml of Isoton II (Beckman Coulter) diluent and 50 μl of this dilution was counted electronically. Three independent growth replicates were performed per mutant. Estimations of the cell growth parameters were performed as per (Rodríguez-Verdugo et al., 2016). Briefly, we estimated the maximum growth rate by fitting a linear regression to the natural logarithm of the cell density over time during the exponential phase. We estimated the final yield from the cell counts at the end of the exponential phase. Three estimates of maximum growth rate and final yield were obtained for each mutant. The exponential phase was confirmed by linear regression (Zwietering et al., 1990). Statistical differences between *rho* and *rpoB* mutants were assessed by two sample *t*-tests. Statistical analyses (fitting of the linear regression to the exponential phase using the *lm* function and *t*-test analyses) were performed in R version 3.0.2 (Team, 2013). Graphs for fig. 2 were constructed with the function stat-smooth in the ggplot2 R package version 3.2.3 (Team, 2013)

### RNA Harvesting and Sequencing

To harvest cells for RNA sequencing (RNAseq), we revived cells as described above and then cultivated populations at 42.2°C until the mid-exponential phase of population growth. The mid-exponential phase was determined by electronic counts using an electronic particle counter (Coulter Counter model Multisizer 3 equipped with a 30 µm diameter aperture tube) and linear regression. For each of the single *rho* mutations, at least two biological replicates were grown to mid-exponential for RNA harvesting.

To harvest RNA, 80 ml of culture from the mid-exponential phase was filtered through a 0.2 µm cellulose membrane (Life Science, Germany). Then cells were washed with Qiagen RNA-protect Bacteria Reagent and pelleted for storage at -80°C prior to RNA extraction. Cell pellets were thawed and incubated with lysozyme (Sigma-Aldrich), for 5 minutes at room temperature ∼20°C. Total RNA was isolated and purified using RNeasy Mini Kit (Qiagen). An on-column DNase treatment was performed for 30 min at room temperature. RNA quality was assessed by running an Agilent RNA-Nano chip on a bioanalyzer. rRNA was depleted using the Ribo-Zero^TM^ rRNA Removal kit for Gram-Negative Bacteria (Epicentre Biotechnologies, Medion, WI, USA). cDNA libraries were constructed using TruSeq RNA v2 kit (Illumina, San Diego, CA, USA). After amplification, the samples were quantified by qPCR using the KAPA Library Quantification Kit code KK4822 for the Illumina Genome Analyzer platform (Kapa Biosystems, Wilmington, MA, USA). Libraries were sequenced on an Illumina Hiseq 2000 platform to generate 100 bp single-end reads.

RNAseq data for *rpoB* mutations and the REL1206 ancestor were generated previously, using identical protocols (Rodríguez-Verdugo et al., 2016).

### RNAseq data and DEG analysis

We gathered three sets of RNAseq data. First, we generated data for four *rho* mutants, with two replicates each. Second, we used the REL1206 data from Rodriguez-Verdugo et al. (2016), representing states of both thermal stress (42.2°C) and normal laboratory conditions (37.0°C). Finally, we also used data from *rpoB* single mutants from (Rodríguez-Verdugo et al., 2016). The *rpoB* data can be accessed at the NCBI’s Sequence Read Archive (SRA) under the BioProject Number PRJNA291128. The *rho* data generated for this study have BioProject Number PRJNA339971 and SRA accession number SRP083065.

For all RNAseq data, sequence reads were filtered to a quality cut-off of 20. Filtered reads were mapped to the *E. coli* B REL606 reference genome (NCBI: NC_012967.1 GI:254160123) using BWA aligner (Li and Durbin, 2010) using default parameters. Statistics about mapping are provided in supplementary table S7 of the Supplementary Material online. Unique matching reads to the 4204 annotated coding regions were retained for further analyses, as were reads that mapped to 2306 non-coding intergenic regions (IRs) of the REL606 reference (supplementary data set S3, Supplementary Material online). IRs were defined as the sequence that extends from the 3’ end of a coding region to its neighboring coding region. IRs were analyzed if they exceeded 1 bp in length and had a minimum coverage of at least 1 read in at least one of the ancestors and *rho* mutants replicates. Note, however, that similar trends were obtained when IRs were defined as ≥100 bp in length.

Coding regions and IRs counts were obtained from SAM files using HTSeq-count tool (Anders et al., 2015) from HTSeq, at http://www.huber.embl.de/users/anders/HTSeq/doc/count.html using intersection-nonempty counting mode. Differential gene expression analysis was performed with DESeq version 1.18.0 (Anders and Huber, 2010). The FDR was determined using Benjamini-Hochberg adjusted *p*-values. Genes with *q* values <0.001 were considered as differentially expressed.

Gene Ontology enrichment analysis was performed through the Gene Ontology Consortium (http://geneontology.org/page/go-enrichment-analysis) with Bonferroni correction for multiple testing. Principal component analysis (PCA) was applied to RNAseq data using the *plotPCA* function in DESeq version 1.18.0 (Anders and Huber, 2010) with the default parameters (

~~~
ntop=500
~~~

).

### Identification and analysis of rho-terminated genome regions

To evaluate the effects of each *rho* mutation on the transcription termination activity, we evaluated the gene expression of a set of previously defined Rho dependent terminated regions (Peters et al., 2009). Because these Rho-terminated regions were originally defined in *E. coli* K-12, it was necessary to first determine their coordinates in our REL606 genome. To do so, we utilized the start positions and lengths of the regions as defined by Peters et al. (2009) located the closest gene to the start of the Rho-terminated region in *E. coli* K-12, noting the position of the Rho-terminated region relative to the closest gene. We then located this gene in the REL606 genome and used the relative position information from K-12 to assign our Rho-terminated regions in REL606. In the few cases where gene orientation was not preserved across strains, coordinates were determined in an orientation-dependent manner (supplementary data set S2, Supplementary Material online). The number of read counts of each region was obtained from SAM files using the HTseq-count tool (Anders et al., 2015) and normalization was performed with DEseq based on a complete gff file that included the putatively Rho-terminated regions and their 5’ upstream region of the same size (Anders and Huber, 2010). Five of 183 regions had no reads and were discarded. Normalized gene counts were averaged among replicates to calculate the ratio *γ*, the log_2_ ratio of mutant to ancestral counts.

For each of the 178 *rho* termination regions, we also compared read counts between the termination region and its immediate upstream 5’ region of the same length. Thus, for each of the *i*=10 treatments (i.e., 8 mutants and 2 control temperatures), the data consisted of paired upstream and downstream counts from *j*=178 regions. For each clone *i*, we estimated termination efficiency by bootstrap resampling across the *j* regions, keeping the 5’ and termination regions paired. For each of 1000 resamplings, we calculated the ratio of averages, *r*_*avg*_, across the *j* samples (fig. 4). To test for differences among a set of *i* treatments, we bootstrapped under the null hypothesis of homogeneity by resampling across the *i* treatments for each of the *j* regions. For each of 1000 datasets resampled under the null hypothesis, we calculated *r*_*avg*_, for each of the *i* treatments. Bootstrap resamplings were performed in R version 3.0.2 (Team, 2013).

### Directions of evolutionary change

GE was classified into one of four directions of evolutionary change, as described previously (Carroll and Marx, 2013, Rodríguez-Verdugo et al., 2016). The four directions classify GE in a mutant relative to REL1206 at 42.2°C and to REL1206 at 37.0°C. Briefly, a gene had *novel* expression if the mutant differed significantly (*q* < 0.001) from both ancestral treatments (42.2°C and 37.0°C), but GE did not differ between the two ancestral treatments (i.e. Anc42≈Anc37 & Anc37≠Mutant & Anc42≠Mutant). *Restoration* occurred when the mutant GE levels fell between those of the two ancestral treatments and when GE differed significantly between the ancestral treatments and also between the mutant and REL1206 at 42.2°C [i.e., (Anc42>Mut>Anc37 or Anc42<Mut<Anc37) & Anc42 ≠Anc37 & Anc42 ≠Mutant]. *Reinforced* expression occurred when GE differed significantly between the ancestral treatments and between the mutant and REL1206 at 42.2°C, but GE in the mutant did not fall between the ancestral treatments [i.e., Anc42 ≠Anc 37 & Anc42 ≠Mutant & (Mut>Anc42>Anc37 or Mut<Anc42<Anc37)]. Finally, *unrestored* expression occurred when GE did not differ significantly between the mutant and REL1206 at 42.2°C, but the two ancestral treatments differed significantly from each other (i.e., Anc42 ≠Anc 37 & Mut≈Anc42).

### Estimates of free energy of folding

We calculated two values: ΔG, the free energy of folding, and ΔΔG, the difference of ΔG between the REL1206 protein and mutant proteins. We focused on folding free energy differences of the Rho hexamer and *rpoB*. To calculate ΔΔG, we used an approach that combines classical molecular dynamics with FoldX (Guerois et al., 2002, Miller et al., 2016), which improves the calculation of ΔΔG compared to using FoldX alone (Miller et al., 2016). To perform the calculations, we downloaded the X-ray crystal structures from the Protein Data Brank (PDB): (1) Rho transcription protein bound to RNA and ADP-BeF3 (PDB ID: 3ICE) (Thomsen and Berger, 2009) and (ii) SigmaS-transcription initiation complex with 4-nt nascent RNA (PDB ID: 5IPL) (Liu et al., 2016). The PDB file 3ICE was modified to remove all but the six chains of Rho hexamer. Out of all the six chains in the X-ray crystal structure, chain C had least missing residues. MODELLER v9.15 software (Sali and Blundell, 1993) was then used to rebuild the coordinates of the missing residues for chain C and complete chain C was used as a template to fill gaps in the other chains of the hexamer. The final hexamer structure had 2484 residues. The chain C of 5IPL PDB file was used to obtain the structure of *rpoB* which had 1342 residues.

The software package GROMACS 5.0.7 (Van Der Spoel et al., 2005) was used for 10 ns long Rho hexamer and *rpoB* Molecular Dynamics simulations with the AMBER99SB forcefield (Hornak et al., 2006). We followed the standard energy minimization, thermalization and equilibration protocol (Miller et al., 2016) before carrying out the final production run. During the 10 ns production simulation snapshots were saved every 1 ns giving 10 snapshots of Rho hexamer protein and *rpoB* for the further analysis.

Fold X was used to analyze the starting structure and all 10 snapshots (total 11 snapshots) captured during molecular dynamics simulations (Guerois et al., 2002). Before, analyzing all 11 snapshots using FoldX, we first extracted chain C from each of the snapshots in case of Rho hexamer and *rpoB* and later only the monomer structures - i.e. chain C - were used for folding free energy calculations. We started by minimizing monomer structure using the *RepairPDB* command of FoldX to further minimize the potential energy. All single amino acid mutations were then generated using *BuildModel*. Finally, protein folding stabilities were estimated using *Stability* on the *rho* monomer and *rpoB* structure. For each mutation we then estimated ΔΔG by averaging across all 11 individual snapshots estimates. The molecular graphics package VMD was used to produce fig. 1(Humphrey et al., 1996).

## ACKNOWLEDGMENTS

We thank R. Gaut and P. MacDonald for technical assistance. A.G.G. and A.R.V. were supported by University of California Institute for Mexico and the United States-Consejo Nacional de Ciencia y Tecnología (Mexico) Post-doctoral and Doctoral fellowships respectively. S.H. was supported by the National Institute of Biomedical Imaging and Bioengineering, National Research Service Award EB009418 from University of California, Irvine, Center for Complex Biological Systems. J.S.P. was supported by the National Institute Of General Medical Sciences of the National Institutes of Health under Award Number P20GM104420 from Center for Modeling Complex Interactions, University of Idaho, Moscow. BSG was supported by a fellowship from the Borchard Foundation, and the work was also supported by the National Science Foundation grant DEB-0748903. Computer resources to perform the Molecular dynamic simulations were provided by the Institute for Bioinformatics and Evolutionary Studies Computational Resources Core sponsored by the National Institute of Health (P30 GM103324) at University of Idaho.

